# Fecal Microbiota Restoration Modulates the Microbiome in Inflammation-Driven Colorectal Cancer

**DOI:** 10.1101/2022.12.28.522045

**Authors:** Travis J Gates, Ce Yuan, Mihir Shetty, Thomas Kaiser, Andrew C Nelson, Aastha Chauhan, Timothy K Starr, Christopher Staley, Subbaya Subramanian

## Abstract

Chronic inflammation of the colon (colitis) is a known risk factor for inflammatory-driven colorectal cancers (idCRC), and intestinal microbiota has been implicated in the etiology of id-CRC. Manipulation of the microbiome is a clinically viable therapeutic approach to limiting id-CRC. To understand the microbiome changes that occur over time in id-CRC, we used a mouse model of idCRC with the treatment of azoxymethane (AOM) and dextran sodium sulfate (DSS) and measured the microbiome over time using 16S rRNA amplicon sequencing. We included cohorts where the microbiome was restored using cage bedding swapping and where the microbiome was depleted using antibiotics to compare to untreated animals. We identified consistent increases in *Akkermansia* in mice receiving horizontal microbiome transfer (HMT), while the control cohort had consistent longitudinal increases in *Anaeroplasma* and *Alistipes*. Additionally, fecal lipocalin-2 (Lcn-2) concentrations were elevated in unrestored animals compared to restored and antibiotic-treated counterparts following HMT. These observations suggest a potential role for *Akkermansia, Anaeroplasma, and Alistipes* in regulating colonic inflammation in id-CRC.

## INTRODUCTION

Colorectal cancer (CRC) is the second leading cause of cancer-related death worldwide.^1^ Inflammation of the colon has been shown to contribute to the development of CRC through various mechanisms, such as changing the local and systemic cytokine milieu through alteration of NF-kB and STAT3 signaling pathways promoting growth and survival.^2,3^ Additionally, colitis has increased the likelihood of induced gene mutations of the colonic epithelium and epigenetic alterations, providing conditions for growth.^4-6^ Clinical observation has also suggested a correlation between colonic neoplasia and increased mucosal prostaglandin, implicating a link between inflammation and CRC progression. Inflammatory or colitis-driven CRC presents a unique microenvironment in which microbial dysbiosis has been implicated in driving or supporting tumor initiation and progression.^7-9^ Dysbiosis of the microbiota can be loosely defined by a reduction in total microbial diversity and reduced abundances of commensal bacteria such as *Firmicutes* and *Bacteroidetes*. These commensal bacteria can provide the host with short-chain fatty acids such as butyrate, which has been shown to suppress colonic inflammation.^10,11^ This is coupled with the increased abundance of pathobionts, such as members of the family *Enterobacteriaceae*.^*12,13*^ Yet, these studies have yet to reach a consensus.

Many factors complicate determining causative associations between the microbiome and CRC pathogenesis. These factors include environmental, inter-individual, and dietary differences, variable experimental conditions such as sampling parameters, i.e., stool or mucosa, longitudinal fluctuations, and lack of batch-to-batch reproducibility.^14-16^ Advantages of using experimental mouse models include their homogenous genetic background, consistent environment and diet, and accessible longitudinal sampling, which can be temporally controlled. The most widely used model to induce colonic inflammation consists of dosing a single *i*.*p*. injection of azoxymethane (AOM) which cause DNA damage and is a carcinogen. This is followed by an infusion of dextran sodium sulfate (DSS) into the mice’s drinking water, which induces colonic inflammation.^17^ This AOM/DSS model induces epithelial injury and promotes CRC development, mimicking human inflammation-driven CRC.^18^

Some studies have reported microbial composition and diversity alterations following AOM/DSS induction.^19-21^ Others have shown that unique fecal compositions can be transplanted into germ-free mice and are associated with different tumor outcomes through altering T-cell responses in AOM/DSS induced models.^22^ Fecal microbiota transplantation (FMT) of a CRC patient’s stool into *APC*^*min/+*^ mice was also shown to lead to the accumulation of more intestinal tumors compared to healthy donor controls, suggesting a role of a tumor microbiome in CRC progression.^23^ Restoration of the fecal microbiota via FMT is an interesting approach to understanding microbial ecology. It is being explored in many colon diseases, including recurrent *Clostridiodes difficile* infection^24^, irritable bowel disease,^25,26^ and CRC.^27^ Further, it has been shown that cohousing of animals can modulate the microbiome between mice through horizontal microbiome transfer (HMT).^28^

The longitudinal effects on the gut microbiota following HMT in AOM/DSS models of CRC have not been explored. In this study, we longitudinally analyzed the fecal microbiome of C57BL/6 and BALB/c mice that underwent swapping of used bedding to achieve HMT and homogenize the microbiome with normal mice (restored) to compare to mice that received no transfer (unrestored). We observed significant and different early to late changes between restored and unrestored groups, which were consistent between C57BL/6 and BALB/c species following AOM/DSS administration.

## RESULTS

### Microbial community composition and diversity between restored and unrestored groups are longitudinally different

We longitudinally compared the microbial community changes following AOM/DSS administration between unrestored (untreated) and restored (HMT from normal Balb/c fecal bedding swap once weekly during weeks 8, 9, and 10). Mouse fecal pellets were longitudinally collected three times weekly for 17 weeks from Balb/c mice in unrestored n=5 and restored n=5 groups (**Figure 1A**). The overall mean percent relative abundance of bacterial genera in both restored and unrestored groups was relatively stable and similar **(Figure 1B)**. Longitudinally, the microbial communities in mice with restored microbiota had significantly lower Shannon index than unrestored mice (p=0.035**; Figure 1C)**; however, no significant differences were observed in Chao1 (p=0.084**; Figure 1D)**.

**Figure 1:**
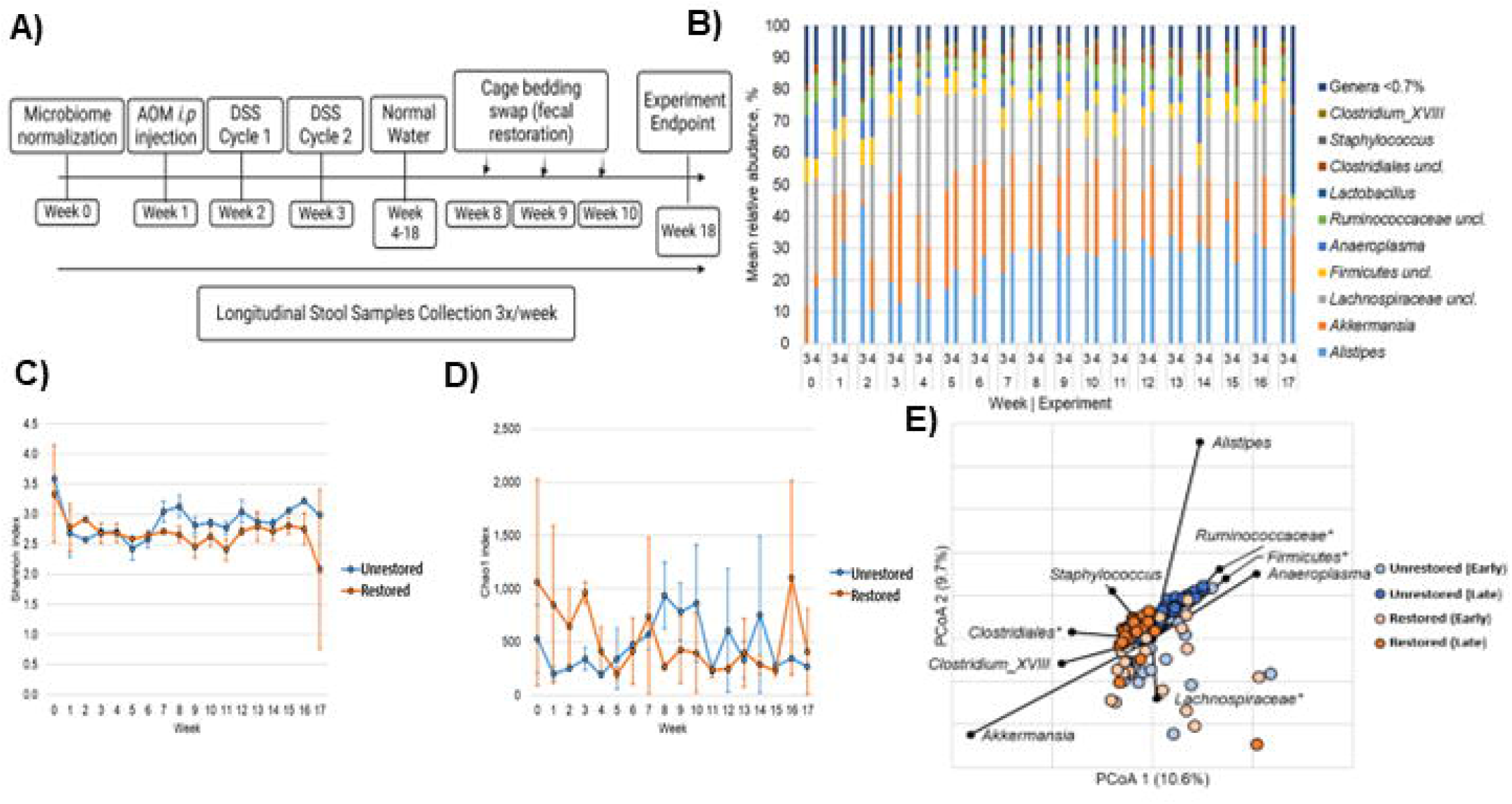
Microbiome composition and diversity in Balb/c mice. **A)** Percent reiative abundance of bacterial genera in Balb/c mice following AOM/DSS administration between (3; unrestored) and (4; restored) groups. Genera that accounted for <0.7% mean sequence reads were consolidated for clarity. **B)** Depicts longitudinal Shannon index between no-restoration (blue) and restoration (orange). Error bars reflect standard deviation. **C)** Depicts longitudinal Chao1 index between no-restoration (blue) and restoration (orange). **D)** Principal coordinates analysis with correlated taxa associated with ordination position on either axis (Spearman p>0.05) are overlaid on the PCoA plot.

Next, we sought to investigate the difference in community composition between restored and unrestored groups using Bray-Curtis dissimilarity. Samples in both treatment groups were considered as “early” (weeks 0-7, before microbiota restoration) and “late” (weeks 8-18; at the time of and following HMT). Principal coordinates analysis (PCoA) showed significant early and late changes in the microbiome of Balb/c mice treated with AOM/DSS **(Figure 1E)**. The unrestored group had significant early and late changes (ANOSIM R=0.499, p<0.001) within the group **(Figure 1E)**. Similarly, we also observed significant early and late changes in the restored group (ANOSIM R=0.261, p<0.001). The microbiota composition also differed significantly between experimental groups among the later “late” samples but not samples collected prior to week 8 (R=0.530 and 0.081, p<0.001 and p=0.01; Bonferroni-corrected α=0.008**; Figure 1E)**

Spearman correlation was used to identify bacterial genera associated with the late differences in the microbial community between unrestored and restored groups (**Figure 1E**). *Akkermansia, Clostridium_*clade XVIII, and members of the order *Clostridiales* that could not be further classified were compositionally correlated with the restored group **(Figure 1E)**. This was an interesting observation as *Akkermansia* and *Clostridium*_XVIII have been implicated as having probiotic properties and associated with reduced intestinal inflammation by producing short-chain fatty acids like butyrate.^40,41^

In contrast, microbial communities in the restored group were correlated with greater relative abundances of the bacterial genera *Alistipes, Anaeroplasma, and Ruminococcaceae* **Figure 1E**. We were unable to further classify *Rumminococcaceae* in our 16S rRNA analysis. Still, interestingly *Ruminococcus gnavus* is associated with Crohn’s disease, likely through *R. gnavus* ability to synthesize and secrete glucorhamnan polysaccharides, which can lead to TNFα secretion by dendritic cells.^42^ Additionally, the increased abundance of potentially pathogenic *Alistipes* is consistent with recent investigations of AOM/DSS induced cancer.^43^ These data suggest that the HMT altered the microbial composition in the restored group compared unrestored group, potentially shifting toward a less inflammatory community composition.

### Longitudinal differences in the microbiome between restored and unrestored groups is significant

Next, we sought to investigate longitudinal changes in bacterial genera between restored and unrestored groups using the SplinectomeR permuspliner function, which can be used to assess longitudinal microbiome data.^39^ Significant longitudinal differences were observed between groups among the genera *Alistipes* (p= 0.032), *Akkermansia* (p=0.001), *Anaeroplasm*a (p=0.032), *Ruminococaceae* (further unclassified) (p=0.001), *Clostridiales* (further unclassified) (p=0.001) *Clostridium_XVIII* (p=0.001) **(Supplemental Figure 1A-F)**. Significant longitudinal differences in Shannon index (p=0.035) between groups were also observed **(Supplemental Figure 1G)**. Notably, the longitudinal significant difference between unrestored and restored is somewhat a conflicting result as previous work has shown that alpha diversity has decreased longitudinally in AOM/DSS models of intestinal inflammation.

The longitudinal abundance of *Akkermansia* was significantly different between unrestored and restored groups following fecal restoration (**Supplemental Figure 1A)**. For *Akkermansia, the* mean relative abundance was significantly greater in the restored group compared to the unrestored in weeks 9-17. This was also reflected in our Bray-Curtis dissimilarity and Spearman correlation analysis in **Figure 1D**. Many bacterial genera identified from Spearman correlations were also longitudinally altered between unrestored and restored in SplinectomeR permuspliner analysis.

### Intestinal inflammation marker Lipocalin-2 is decreased in restored Balb/c mice

To assess differences in intestinal inflammation before and after HMT in AOM/DSS treated animals, we utilized a semi-quantitative sandwich ELISA (R&D systems, Minneapolis, MN) to probe fecal concentrations of Lipocalin-2 (Lcn-2).^44,45^ Fecal Lcn-2 concentrations are a non-invasive marker of intestinal inflammation. Two time points were used before HMT (weeks 2 and 7), and two-time points post HMT (weeks 10 and 15) were selected to quantify fecal concentrations of Lcn-2 in restored and unrestored groups. In weeks 2 and 7, we did not observe significant differences in fecal Lcn-2 concentrations (Week 2, 136.1 ± 8.0 pg/mL and 123.7 ± 9.8 pg/mL p=0.082, n=3) (Week 7, 171.1 ± 21.7pg/mL and 154.2 ± 17.2pg/mL p=0.154, n=3) between unrestored and restored animals respectively **Figure 2A**. Post HMT, we did observe significant decreases in fecal Lcn-2 concentrations (Week 10, 206.1 ± 20.9 pg/mL and 139.4 ± 17.8pg/mL p=0.007, n=3) (Week 15, 335.4 ± 16.2 pg/mL and 252.4 ± 27.3 pg/mL p=0.005, n=3) in unrestored and restored animals respectively **Figure 2A**. These data suggest that HMT could potentially reduce Lcn-2 concentrations and could have transiently reduced intestinal inflammation in restored animals compared to unrestored counterparts.

**Figure 2:**
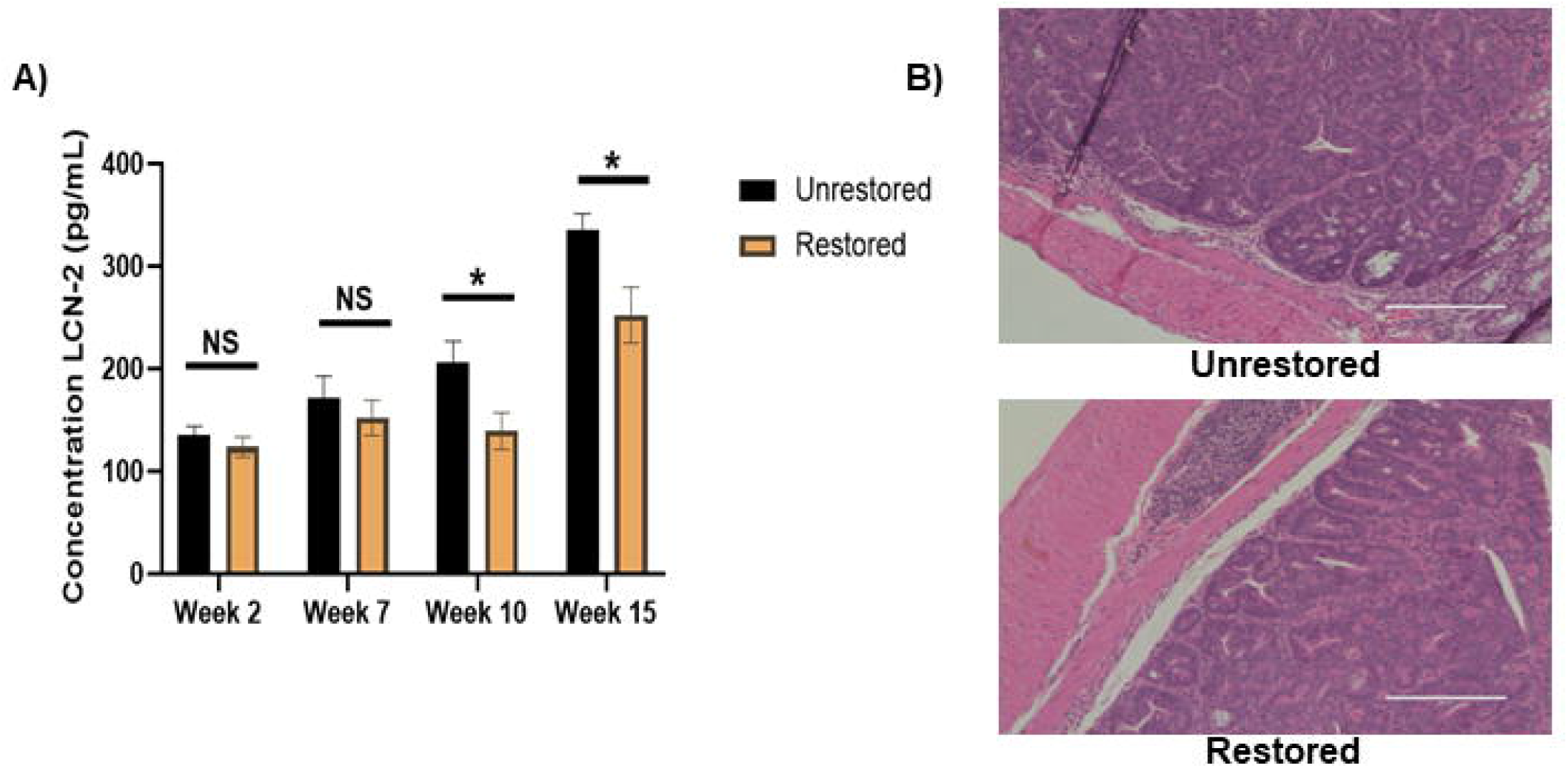
Fecal restoration alters colonic inflammation in Balb/c mice. **A)** Fecal concentrations of Lcn-2 in unrestored (black) and restored (orange) in weeks 2, 7, 10,15 (left to right) (*p<0.05) (n=3) (+/- SD) **B**) Hematoxylin and eosin staining of colon tissues of unrestored and restored Balb/c mice.

Despite significant differences in fecal Lcn-2 concentrations, we did not observe any pathological differences at the experiment endpoint when looking at the hematoxylin and eosin staining of fixed colon tissues **Figure 2B**. Both restored and unrestored animals at the experiment endpoint presented histology ranging from high-grade dysplasia to high-grade intramucosal adenocarcinoma. Additionally, restored and unrestored animals presented with acute and chronic inflammation ranging from focally mild to focally moderate and focally marked. These observations also align with attempted colonoscopy of unrestored and restored animals in the early stage. However, a colonoscopy was impossible at late stages due to bloody stool and inflammation within the colon. Although there were no significant histological differences at the experiment endpoint, we observed significant differences post-HMT in fecal Lcn-2 concentrations, which led us to believe that the protective effects of HMT were likely transient and the more frequent treatment with HMT could impact the pathology between groups.

### Bacterial community compositions are similar between Balb/c and C57BL/6 in late-restored groups

We previously observed altered longitudinal fecal microbiomes of restored Balb/c compared to unrestored mice. Considering the variable fluctuations in microbiome analysis, we questioned whether our observations were specific to Balb/c. We used 3-4 week-old Balb/c and C57BL/6 mice in the AOM/DSS model to test this. Additionally, we included a group of 3-4 week C57BL/6 mice treated with an antibiotics cocktail of vancomycin 0.05g/L, metronidazole 0.025g/L, streptomycin 0.2g/L to deplete mouse microbiomes as a control, to ensure our longitudinal data acquisition and analysis pipelines were working appropriately. Interestingly, antibiotics-treated mice observed lower-grade dysplasia compared to unrestored and restored counterparts, which was in line with previous literature.^20^ Fecal restoration via HMT was again performed during weeks 8, 9, and 10

Longitudinally stable mean relative abundances of genera between groups Group 6A (Balb/c), Group 6B (C57BL/6 unrestored), Group 7 (C57BL/6 +Antibiotics), and Group 8 (C57BL/6 restored) were observed **(Figures 3A and B)**. These data from C57BL/6 were consistent with our previous observation in Balb/c mice. As expected, we observed depleted alpha diversity in Shannon and Chao1 indices for mice receiving antibiotics compared to restored and unrestored **Figure 3C and Supplemental Figure 2**. Similar alpha diversity was observed between the restored and unrestored groups, which were not significant when compared using ANOSIM. Interestingly, we did not notice decreasing alpha diversity in restored C57BL/6 mice compared to unrestored, as was observed in Balb/c mice.

**Figure 3:**
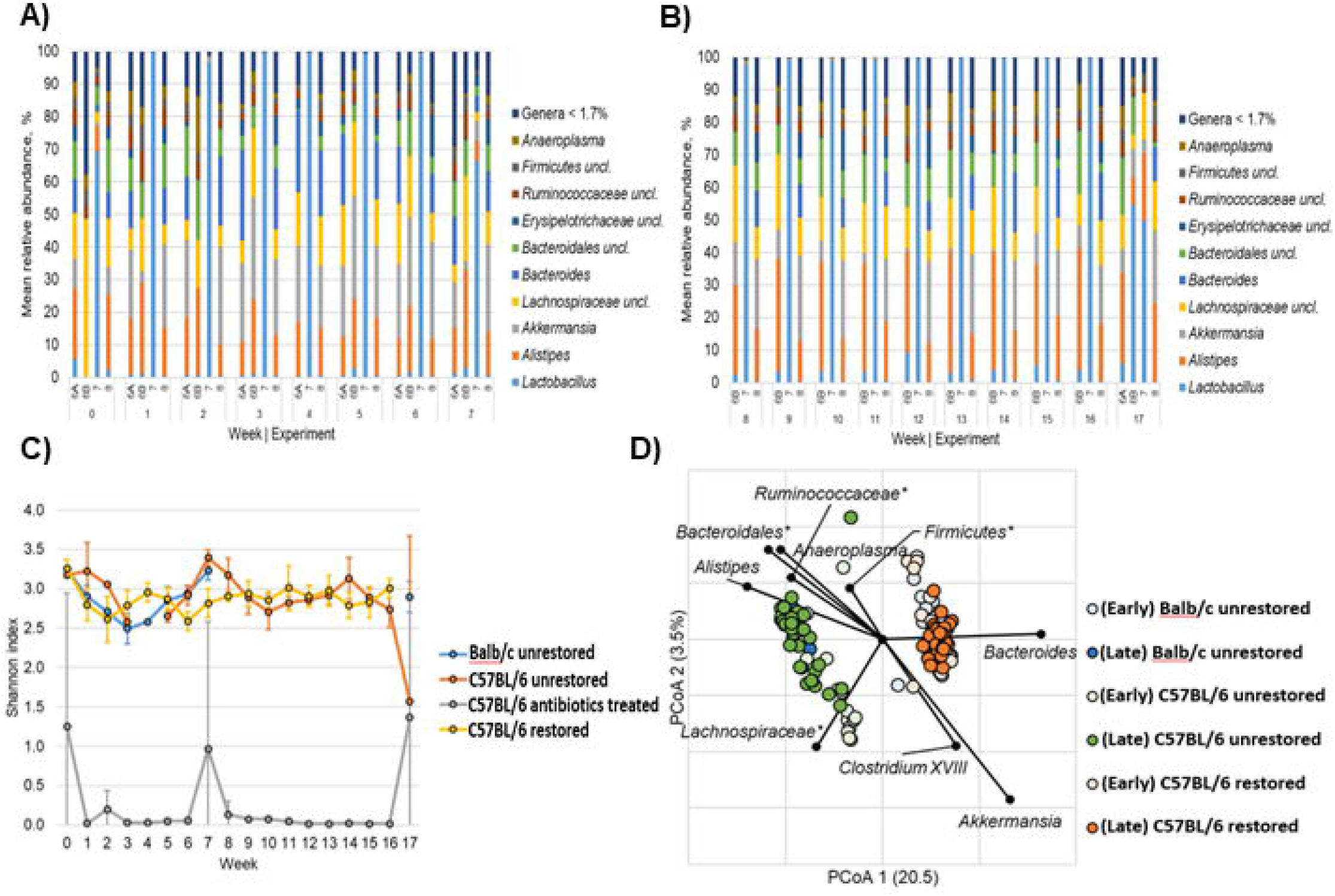
Microbiome composition and diversity in C57BL/6 mice. **A)** Percent relative abundance of bacterial genera (W0-W7) in EXP6A (Balb/c unrestored) EXP6B (C57BL/6 unrestored) EXP7 (C57BL/6 + ABX) and EXP8 (C57BL/6 C57BL/6 restored) following AOM/DSS administration. Genera that accounted for <0.7% mean sequence reads were consolidated for clarity. **B**) Percent relative abundance (W8-17) **C**) Depicts longitudinal average Shannon index between blue (unrestored Balb/c), orange (unrestored C57BL/6), yellow (restored C57BL/6), gray (antibiotics C57BL/6). **D**) Principal coordinates analysis with correlated taxa associated with ordination position on either axis (Spearman p>0.05) are overlaid on the PCoA plot.

Next, we used Bray-Curtis dissimilarity and PCoA to compare restored and unrestored groups’ early and late microbial compositions. The early compositional signature between Balb/c and C57BL/6 unrestored animals was significantly different (ANOSIM R=0.833, p<0.001**; Figure 3D)**. However, unrestored mice showed more similarity in the late microbial compositions between Balb/c and C57BL/6 (ANOSIM R=-0.101; p=0.723; **Figure 3D**). These observations suggested that even though C57BL/6 and Balb/c unrestored mice had significantly different initial microbial compositions after AOM/DSS treatment, similar compositions were observed between strains of mice when left untreated.

Furthermore, we also observed significant differences between late restored C57BL/6 and unrestored C57BL/6 (R=0.897, p<0.001**; Figure 3D)**. Spearman correlation was used to investigate bacterial genera associated with differences in ordination position. Interestingly, we noticed similar taxa (i.e., *Bacteroides, Clostridium_*XVIII, and *Akkermansia)* were correlated with late compositions of the restored group (**Figure 3D)**, similar to our prior observations in Balb/c mice. Additionally, unrestored C57BL/6 communities were associated with a greater relative abundance of *Anaeroplasma, Alistipes, Lachnospiraceae, Bacteroidales* (not further classified), and *Ruminococcaceae* (not further classified). Notably, the genera *Ruminococcaceae, Alistipes*, and *Anaeroplasma*, were found in greater abundance in late samples among untreated groups of both Balb/c and C57BL/6 mice. Similarly, *Akkermansia* and *Clostridium_*XVIII were found at greater relative abundances in the restored microbiota of both mouse genotypes.

### Inflammatory and pathological signatures between Balb/c mice and C57BL/6 mice are different

We again used ELISA to probe fecal concentrations of Lcn-2 pre-HMT (weeks 2 and 7) and post-HMT (weeks 10 and 15) in restored, unrestored, and antibiotics treated animals to observe alterations in intestinal inflammation between groups. We did not observe significant differences in fecal Lcn-2 concentrations pre-HMT (week 2, 115.3 ± 27.4pg/mL and 110.1 ± 19.3 pg/mL p=0.400, n=3) (Week 7, 178.6 ± 26.0pg/mL and 208.6 ±35.1pg/mL p=0.150, n=3) between unrestored and restored animals respectively **Figure 4A**. We did observe significant differences in fecal Lcn-2 concentrations post-HMT (week 10, 200.8 ± 15.3pg/mL and 131.4 pg/mL ± 24.5 pg/mL p=0.007, n=3) between unrestored and restored groups, respectively **Figure 4A**. This result was consistent with our previous investigation in Balb/c mice post-HMT. However, in week 15 we did observe a difference in fecal concentration of Lcn-2 (188.8 ± 29.5 pg/mL and 152.5 ± 15.1pg/mL p=0.065, n=3) between unrestored and restored, but it did not reach statistical significance p=0.065. Interestingly, in all four time points, antibiotics-treated animals had statistically significantly lower concentrations of fecal Lcn-2 (44.2 ± 9.7pg/mL, p=0.003, n=3) (83.5 ± 18.9pg/mL, p=0.003, n=3) (78.1 ± 22.1pg/mL, p=0.024, n=3) (97.9 ± 23.8pg/mL, p=0.014, n=3) than both unrestored and restored animals **Figure 4A**. This was in line with previous investigations in C57BL/6 animals treated with antibiotics in AOM/DSS models of intestinal inflammation. Additionally, in general, Balb/c mice, compared to C57BL/6 mice, presented with elevated levels of fecal Lcn-2 across all treatment conditions, which was also recapitulated in H and E stains of fixed colon tissues in **Figure 2**.

**Figure 4:**
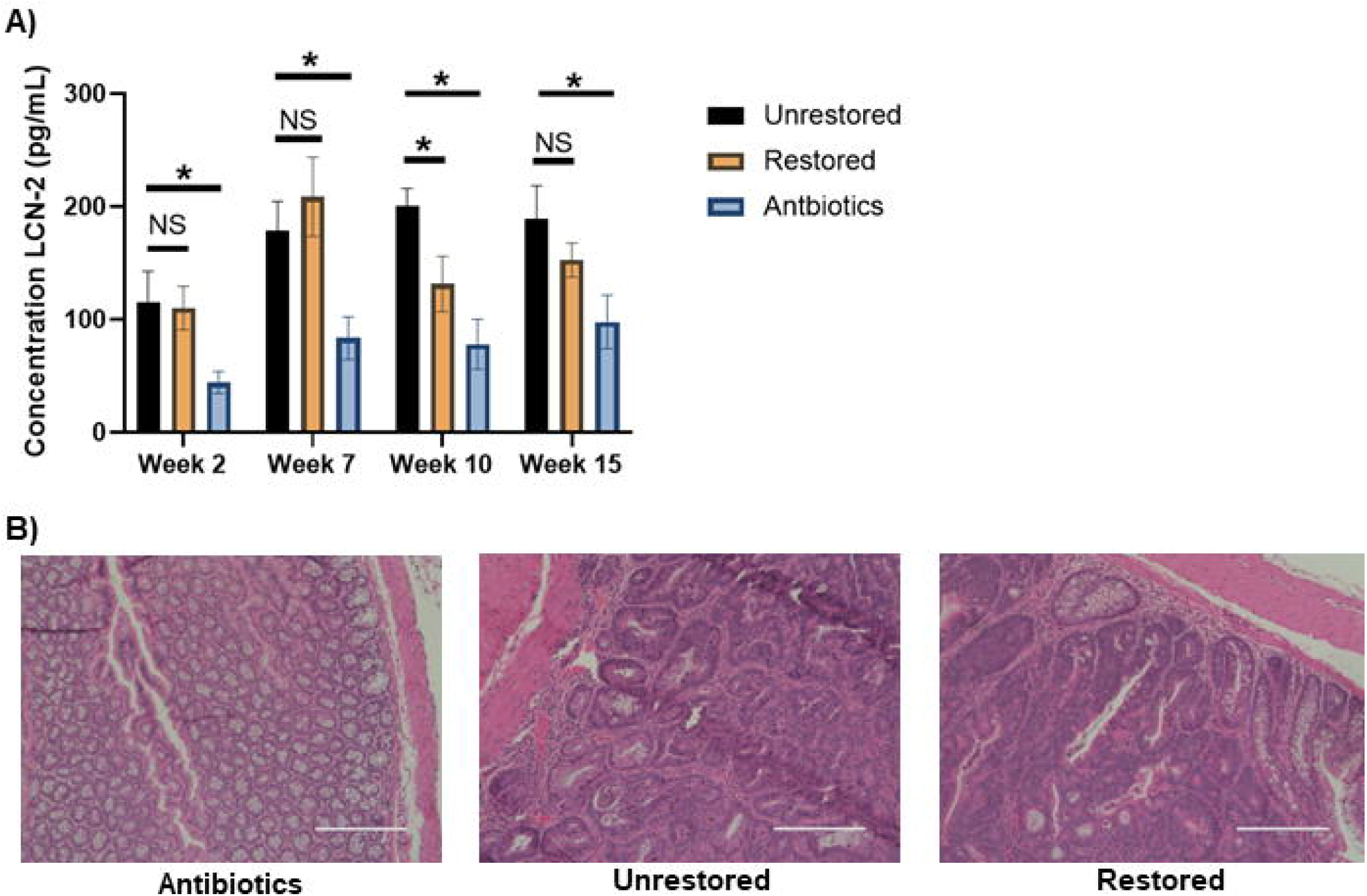
Fecal restoration modulates intestinal inflammation in C57BL/6 mice. **A) C)** Fecal concentrations of Lcn-2 in unrestored (black) restored (orange) and antibiotics treated mice (blue) in weeks 2,7,10,15 (left to right) (*p<0.05) (n=3) (+/- SD) **B**) Hematoxylin and eosin staining of colon tissues of unrestored, antibiotics treated, and restored C57BL/6 mice respectively at 20x amplification.

Even though we observed a statistically significant decrease in fecal Lcn-2 concentrations in week 10 following HMT, this did not influence the pathological observations at the experimental endpoint between restored and unrestored animals **Figure 4B**. Mice in restored and unrestored groups at the experiment endpoint presented with dysplasia ranging from high-grade dysplasia to high-grade intramucosal adenocarcinoma. Furthermore, both restored and unrestored animals presented with acute inflammation ranging from minimal to focally moderate and chronic inflammation from mild to focally moderate. These observations suggested that any beneficial effect from HMT, as determined by concentrations of fecal Lcn-2, was likely transient and did not influence overall pathological outcomes between unrestored and restored groups. However, mice receiving antibiotics were absent of dysplasia and showed only minimal acute-and mild chronic -inflammation. This was consistent with fecal Lcn-2 concentrations as throughout all sampled time points, antibiotics-treated animals had significantly reduced levels of Lcn-2 compared to restored and unrestored animals. These data further support a role or niche of the gut microbiota in regulating intestinal inflammation as mice treated with antibiotics presented with less advanced stage disease compared to unrestored and restored groups.

### *Alistipes, Akkermansia, and Anaeroplasma* are longitudinally altered in C57BL/6 restored and C57BL/6 unrestored mice

Next, we investigated longitudinal changes in bacterial genera between C57BL/6 restored and unrestored groups using the SplinectomeR permuspliner function.^39^ We observed significant longitudinal differences in *Alistipes* (p= 0.001), *Akkermansia* (p=0.001), *Bacteroides* (p=0.001), *Erysipelotrichaeae* (p=0.001), and *Anaeroplasma* (p=0.001) between unrestored and restored groups (**Figure 5 and Supplemental Figure 3)**. These bacterial genera were also identified in our Bray-Curtis dissimilarity analysis and correlated using Spearman correlation. Additionally, we did see longitudinally significant differences in *Bacteroides* between unrestored C57BL/6 mice and restored C57BL/6 mice. However, this was not observed in Balb/c. *Compared to healthy controls, Bacteroides are commonly observed in patients with irritable bowel disease*.^46,47^ When comparing Balb/c and C57BL/6 from all experiments, we consistently saw longitudinal increases in *Akkermansia* in the restored group compared to the unrestored. Along with consistent observations of increased abundance of *Alitipes* and *Anaeroplasma* in unrestored compared to restored in Balb/c and C57BL/6 mice.

**Figure 5:**
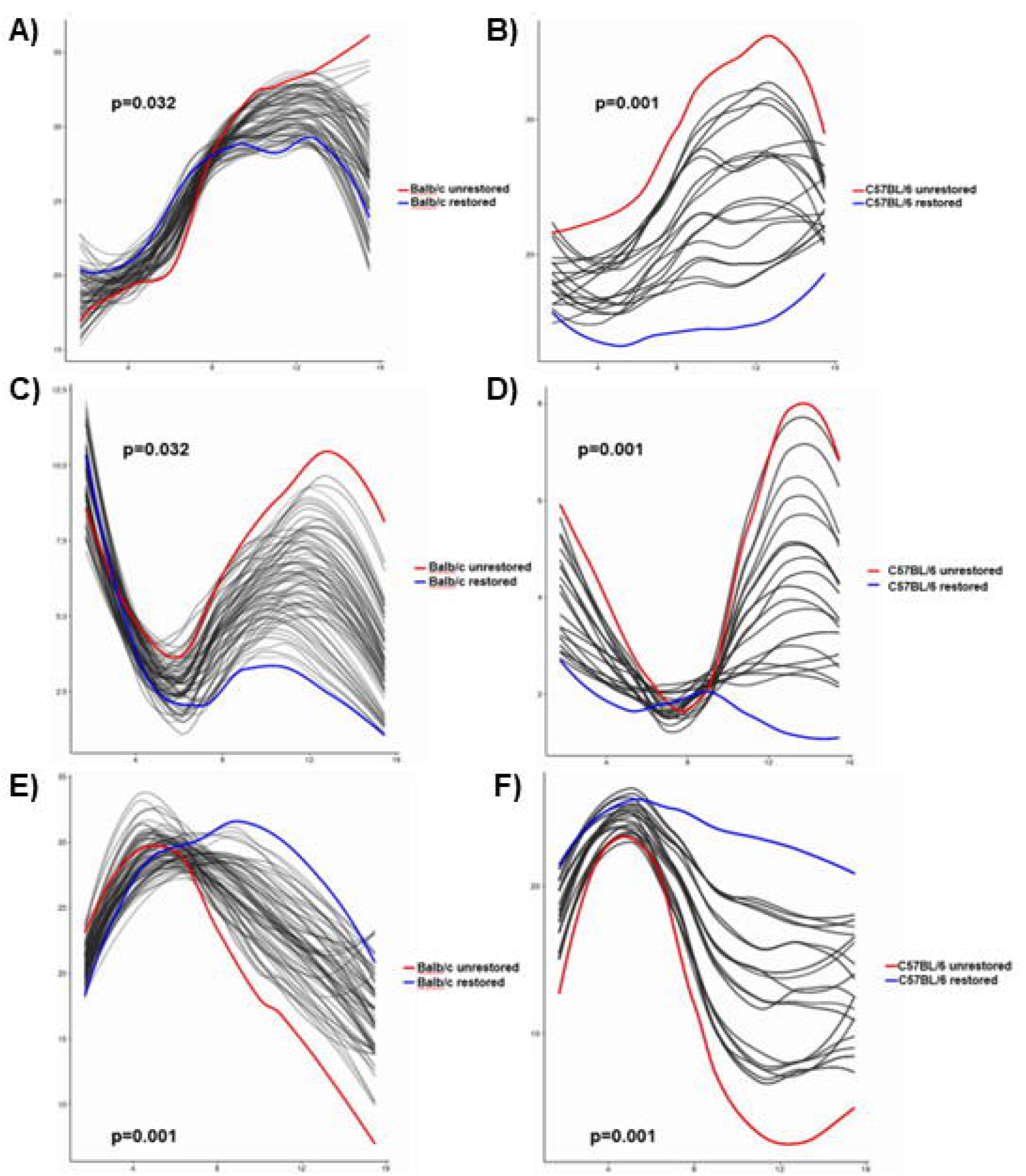
Longitudinally significant genera conserved between mouse genotypes: Depicts longitudinal differences in the relative abundance of genera that significantly differ between (no restoration) red and (restoration) blue determined using SplinectomeR. **A)** *Alistipes*, Balb/c (p= 0.032) **B)** *Alistipes*, C57BL/6 (p=0.001) **C)** *Anaeroplasma*, Balb/c (p=0.032) **D)** *Anaeroplasma*, C57BL/6 (p=0.001) **E)** *Akkermansia*, Balb/c (p=0.001) **F)** *Akkermansia*, C57BL/6 (p=0.001)

## DISCUSSION

Interactions between the microbiota that contribute to host intestinal inflammation are increasingly investigated. Associations have been reported between bacterial genera and disease phenotypes in inflammatory bowel diseases and CRC.^7,48,49^ Many variables render causative associations between the microbiome and disease pathology challenging.^14-16^ The purpose of our study was to use an established mouse model of id-CRC ^17,18^ to longitudinally track shifts in the microbiome of AOM/DSS-treated mice and to assess how fecal restoration influences longitudinal microbial signature and disease pathology. Our study demonstrated that longitudinal compositional changes over 17 weeks are similar between Balb/c and C57BL/6 mice following AOM/DSS treatments **Figures 1 and 3**. Additionally, we have demonstrated that horizontal microbiome transfer (HMT) restoration from untreated mice through bedding swap led to consistent longitudinal increases in *Akkermansia* in Balb/c and C57BL/6 mice compared to their respective unrestored controls. Along with consistent observations of an increased abundance of *Anaeroplasma* and *Alistipes* in unrestored Balb/c and C57BL/6 mice.

Clinical investigations have reported that the relative abundance of *Akkermansia* is decreased in stool samples of patients with active ulcerative colitis compared to patients with quiescent ulcerative colitis and healthy controls.^50^ We reason that observing consistent longitudinal increases in *Akkermansia* in restored mice compared to unrestored mice may represent a more quiescent phenotype following fecal restoration. Additionally, it has been reported that oral administration of *A. muciniphila* strain BAA-835 significantly ameliorated intestinal inflammation following DSS-induced inflammation and was dependent on NLRP3 expression.^51^ Others have shown that *A. muciniphila* strain BAA-835 can reduce inflammatory cytokine expression and reduce mucosal inflammation in DSS models of inflammation.^40,39^ The ability of *Akkermansia* to reduce inflammatory cytokine expression could explain why we observed decreased fecal Lcn-2 in week 10 following HMT in both genotypes of mice **Figures 2 and 4**. Nevertheless, these temporal increases in *Akkermansia* in restored groups did not yield a difference in pathological outcomes between restored and unrestored groups, so this effect was likely transient. More frequent administration of HMT could potentially yield alterations in pathological outcomes, but without further investigation, the overall impact of HMT on disease pathology remains unknown.

Additionally, mice treated with antibiotics observed less dysplasia and inflammation than restored and unrestored animals. Our observations corroborate previous investigations and support that microbiota regulates intestinal inflammation as mice lacking a microbiome observe less aggressive disease.^20^ Furthermore, our Lcn-2 data supports different pathological outcomes in antibiotics treated mice **Figures 2 and 4**. Notably, we also observed significantly lower concentrations of Lcn-2 in C57BL/6 mice compared to Balb/c mice across treatment conditions. This supports our observations from hematoxylin and eosin staining, where Balb/c mice tended to present more frequently with intramucosal adenocarcinoma and focally marked inflammation than C57BL/6 counterparts. In general, C57BL/6 mice presented with less aggressive disease than Balb/c mice. Our studies suggest that these observations should be considered in future investigations of idCRCs, as choosing an appropriate model can influence pathological outcomes. Still, these observations suggest that fecal restoration could contribute to an altered colonic inflammatory response through longitudinal increases in *Akkermansia*.

We also observed significant longitudinal increases in *Alistipes* and *Anaeroplasma* in the unrestored compared to the restored group in Balb/c and C57BL/6 mice **Figures 1, 3 and supplemental Figures 1 and 3**. The observation of an increased abundance of potentially pathogenic *Alistipes* is consistent with recent investigations of AOM/DSS-induced CRCs.^43^ Additionally, more mechanistic insight has been gained by studies showing that *Alistipes* abundance is sufficient to induce colitis in *IL10* ^-/-^ mice and can lead to tumorigenesis.^52^ *Alistipes* has also been identified as a predominant genus of CRC patients from metagenome-wide association studies and is more abundant in carcinomas compared to adenomas and healthy controls.^53^ Our observation of an increased relative abundance of *Alistipes* in unrestored groups compared to restored groups could reflect an increased abundance of *Alistipes* contributing to id-CRCs.

We also observed consistent increases in the relative abundance of *Anaeroplasma* in unrestored compared to restored groups on both murine backgrounds. It has been reported that *Anaeroplasma* is enriched in mice receiving AOM/DSS compared to healthy controls ^21^, and our work demonstrates a similar profile. However, conflicting results have also been reported. For instance, *Anaeroplasma* was shown to be decreased in the stool of mice with common CRC driver gene *APC* mutation compared to the wild type.^54^ However, different inflammatory niches between models could explain this observed difference in the abundance of *Anaeroplasma*.

Further, it has been reported that *Anaeroplasma* strongly correlates with intestinal IgA and TGF-B secretion and regulates intestinal inflammation.^55^ This is also conflicting as IgA increases in patients with IBD^56^, while TGF-B is a known immunosuppressive cytokine.^57^

Lastly, we also observed significant longitudinal increases in *Ruminococcaceae* in Balb/c restored mice compared to unrestored Balb/c but not C57BL/6 mice. Interestingly, *Ruminococcus gnavus* is associated with Crohn’s disease, likely through *R. gnavus* ability to synthesize and secrete glucorhamnan polysaccharides, which can lead to TNFα secretion by dendritic cells^42^ and is enriched in CRC patients compared to normal controls.^58^ We did not see this recapitulated in Balb/c and C57BL/6 mice. However, this suggests that fecal restoration longitudinally shifted the microbiome and modulated fecal Lcn-2 concentrations between restored and unrestored mice between genotypes.

## Conclusion

This study sought to investigate how HMT through cage bedding swapping with untreated animals could influence the microbiome and pathological outcomes of Balb/c and C57BL/6 mice treated with AOM/DSS to mimic id-CRC. We observed consistent and significant increases in *Akkermansia* in restored groups compared to unrestored counterparts in both genotypes. At the same time, we observed consistent increases in *Anaeroplasma* and *Alistipes* in unrestored animals compared to restored counterparts across genotypes. These observations were correlated with reduced fecal Lcn-2 concentrations post-HMT in restored mice compared to unrestored counterparts. We also observed differences in dysplasia and inflammatory signatures between Balb/c and C57BL/6 mice, which should be considered in future investigations. In comparison, fecal restoration through HMT did not yield pathologically different outcomes between restored and unrestored animals. Temporal differences in fecal Lcn-2 concentrations were observed post-HMT. Future studies should seek to illuminate the direct microbiota-host mechanisms of *Akkermansia, Anaeroplasma*, and *Alistipes* for their roles in regulating intestinal inflammation and CRC development in different model systems.

## MATERIALS AND METHODS

### Mice and Animal Husbandry

3–4-week-old female C57BL/6 and BALB/c mice purchased from Jackson Laboratory were used in this study. On arrival at the animal facility, mice were randomly assigned to restored or unrestored groups (n=5/group) and received an ear punch. Bedding on days 1-3 was pooled between experimental groups to normalize the baseline microbiome between restored and unrestored groups. Following baseline normalization, mice underwent AOM/DSS treatment, and stool samples were longitudinally collected weekly for 18 weeks. In week 18, mice were sacrificed and dissected. Mouse colons were fixed in 10% neutral buffered formalin and processed for histological analysis. All mice were housed in Specific Pathogen Free conditions in fully autoclaved cages for the duration of the experiment. The Institutional Animal Care and Use Committee approved all animal studies.

### AOM/DSS Tumor Induction

Depending on experimental design, female 3–4-week-old BALB/c or C57BL/6 mice were injected intraperitoneally with 10mg azoxymethane (Sigma) per kg mouse weight. After five days, mice were treated with two cycles (1 week/cycle) of 2% dextran sodium sulfate (MP Biomedicals M.W.= 36,000-50,000)) given for five days in the drinking water, followed by regular water for the duration of the experiment. Mice were sacrificed in week 18 following AOM induction. Depending on the experimental setup, the antibiotics vancomycin (Boynton Health) 0.05g/L, metronidazole (Boynton Health) 0.025g/L, and streptomycin (Sigma) 0.2g/L cocktail were administered in the drinking water of mice following the second cycle of DSS administration. During necropsy, colons were dissected and flushed with PBS to visualize tumors in the colon. Following visualization of tumors, tumor tissues were fixed in 10% neutral buffered formalin and sectioned.

### Fecal Restoration and Stool Sample Collection

Mouse stool samples were longitudinally collected 3 times weekly for 18 weeks. Stool sample collection went as follows: Mice were separated into individual autoclaved SPF cages lacking food and bedding and left for 1 hour. On average, 3-5 fecal pellets were collected per mouse. Mice were removed from individual cages and returned to their respective cages depending on group assignment. Stool samples were pooled by experimental groups and stored at -80C until processing for DNA isolation. During weeks 8, 9, and 10, the bedding from normal mice without AOM/DSS was distributed to cages for fecal restoration by horizontal microbiome transfer (HMT).

### DNA Isolation and 16S rRNA Amplicon Sequencing and Analysis

Mouse fecal pellets of approximately 0.1g were homogenized and processed using the DNA PowerSoil kit (Qiagen) with the QIACube automated platform following the manufacturer’s inhibitor removal technology (IRT) protocol. The V5-V6 hypervariable regions were amplified and sequenced using the BSF784/1064R primer set.^29^ Sterile water and no template controls were carried through amplification and sequencing and did not produce amplicons. Samples were paired-end sequenced on a single run of a MiSeq platform (Illumina, Inc., San Diego, CA, USA) at a length of 301 nucleotides (nt). All sequence processing was done using Mothur software version 1.35.1.^30^. Raw fastq files were trimmed to 150nt sequences to remove low-quality regions and were paired-end merged using the fastq-join function.^31^ Reads were further trimmed to remove reads with mean quality scores <35 over a 50-nt window, homopolymers >8nt, ambiguous bases, and more that 2 nt mismatches to primer sequences. High-quality sequences were aligned to the SILVA database (version 132)^32^ and subjected to a 2% pre-clustering step to remove likely errors.^33^ Chimeric sequences were identified and removed using the UCHIME package.^34^ Operational Taxonomic Units (OTUs) were assigned at 99% similarity using the complete-linkage clustering algorithm, and taxonomy was assigned using version 16 from the Ribosomal Database Project.^35^ Raw .fastq files are available at https://www.ncbi.nlm.nih.gov/sra/PRJNA915052.

### Histopathological analysis

Mouse colons were fixed overnight in 10% neutral buffered formalin at the experimental endpoint. Following fixation, solutions were changed to 70% ethanol before submission to the clinical and translational sciences institute at the University of Minnesota. Tissues were paraffin-embedded, and two sections per mouse were stained with Hematoxylin and Eosin. Board-certified pathologists reviewed tissue sections for dysplasia and acute and chronic inflammation.

### Quantification of Fecal Lipocalin-2 (Lcn-2) by ELISA

0.2g of frozen fecal samples were reconstituted in PBS with 0.1% Tween 20 (100mg/ml). 3mm glass beads were added to the mixture and vortexed 3 times in 30-second bursts to obtain a homogenous fecal suspension. The samples were then centrifuged for 10 minutes at 12,000rpm at 4°C. Clear supernatants were collected and stored at -20°C prior to analysis. Lcn-2 levels were approximated using a Duoset murine Lcn-2 ELISA kit (R&D Systems, Minneapolis, MN) following the manufacturer’s protocol. 4 time points before and after fecal restoration (W2, W7, W10, W15) were selected for fecal Lcn-2 analysis. Samples for each experimental condition were run in triplicate. Calibration standard curves (500pg/mL-7.81pg/mL) were used to extrapolate Lcn-2 concentrations and averaged across experimental time points and conditions. Reagent blank 1.0% BSA in PBS was used as a negative control.

### Statistics

Alpha diversity of microbial communities was assessed using Shannon and Chao1 indices. Bray-Curtis dissimilarity matrices^36^ were calculated and used for ordination by principal coordinates analysis.^37^ These matrices were also used to determine differences in beta diversity by analysis of similarity (ANOSIM).^38^ Bonferroni corrections for multiple comparisons were performed for ANOSIM analyses. The SplinectomeR, R package,^39^ permuspliner functions were used to assess significant differences in the longitudinal abundances of bacterial genera between restored and unrestored groups using standard 999 permutations. 1-way unpaired equal variance students T-test was conducted to assess statistically significant differences in concentrations of fecal Lcn-2. All statistics were evaluated at α=0.05 unless otherwise corrected for multiple comparisons.

## Acknowledgments

We thank the Masonic Cancer Center’s core facilities, the University of Minnesota Genomics Center, and the Clinical and translational sciences institute for microbiome sequencing and supporting histological services. This study is supported by research grants from the Masonic Cancer Center ChainBreaker Fund, Mezin Koats colorectal cancer research fund, Minnesota Colorectal Cancer Funds, and research funds from the Department of Surgery and CTSI, University of Minnesota. The Minnesota Colorectal Cancer Research Foundation supports TJG graduate fellowship.

## Conflict of Interest

The authors declare no conflicts of interest.

## SUPPLEMENTAL FIGURES

**Supplementary Figure 1:**
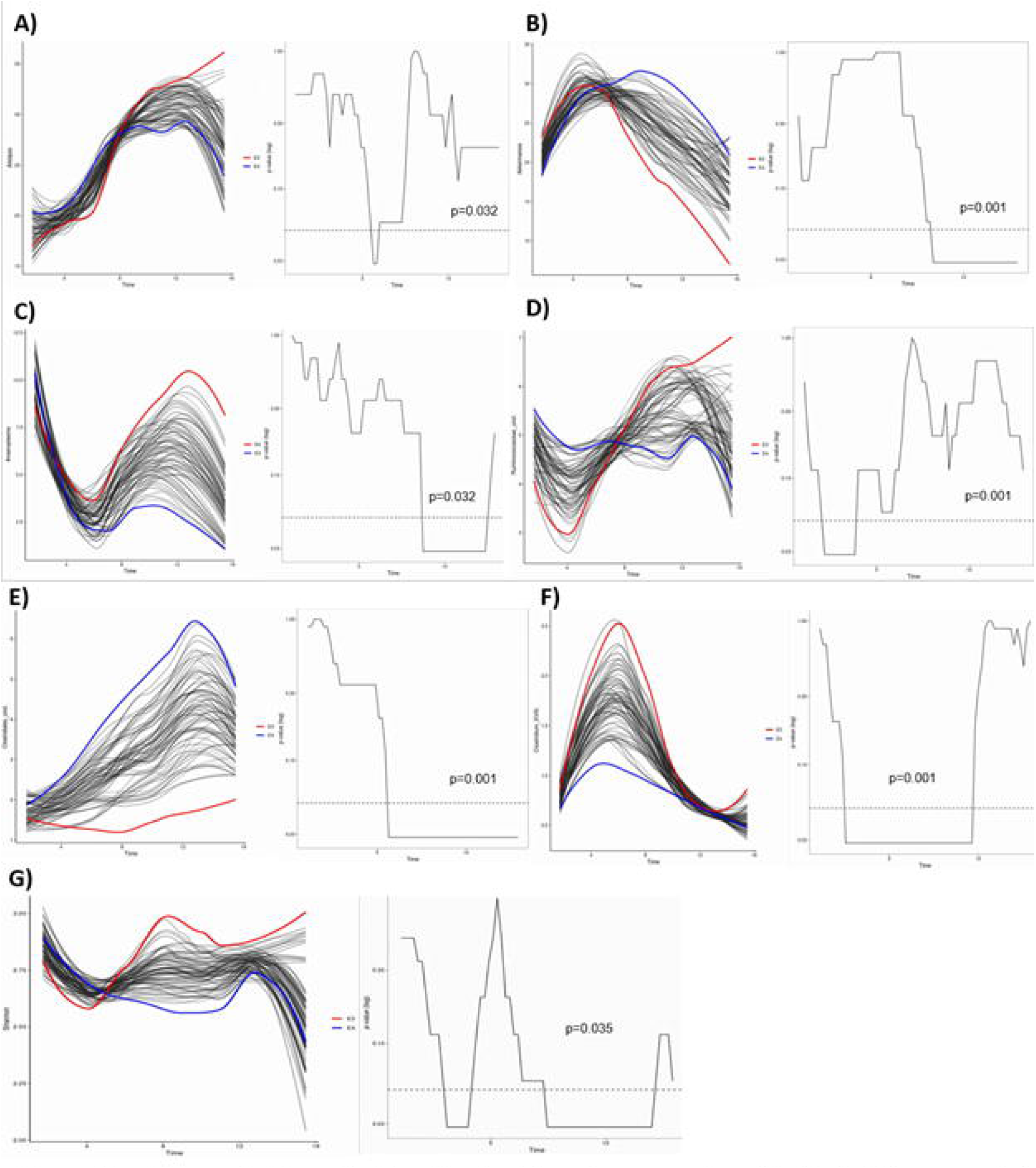
Longitudinally significant genera: Longitudinal differences in the relative abundance of genera that significantly differ between (no restoration) red and (restoration) blue determined using SplinectomeR. **A)** Alistipes (p= 0.032) **B)** Akkermansia (p=0.001) **C)** Anaeroplasma (p=0.032) **D)** Ruminococacea_uncl (p=0.001) **E)** Clostridiales_uncl (p=0.001) **F)** Clostridium_XVIII (p=0.001) **G)** Shannon index (p=0.035)

**Supplemental Figure 2:**
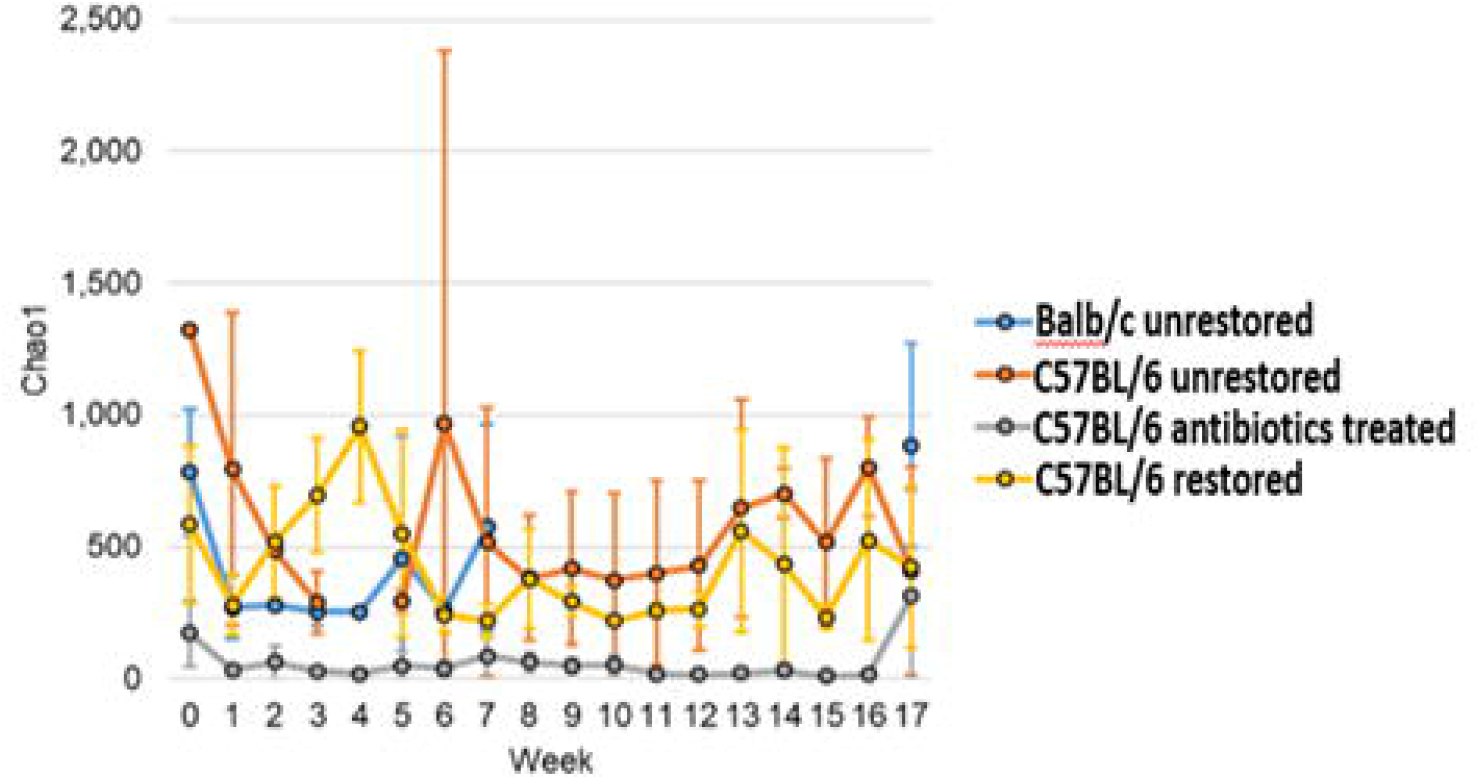
Alpha diversity analysis: Longitudinal average Chao 1 index between blue (unrestored Balb/c), orange (unrestored C57BL/6), yellow (restored C57BL/6), gray (antibiotics C57BL/6).

**Supplemental Figure 3:**
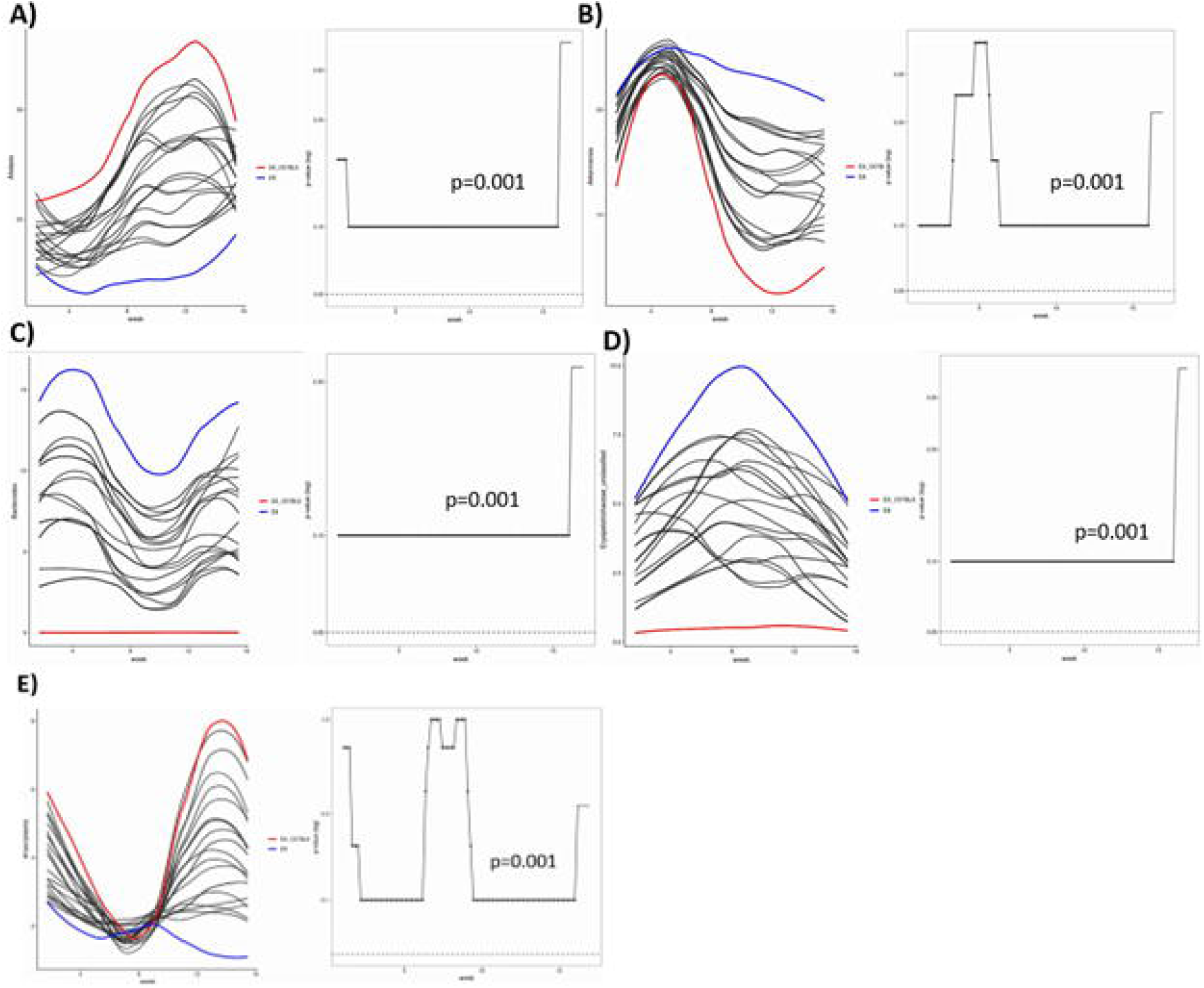
Longitudinally significant genera: Longitudinal analysis between unrestored C57BL/6 (red) and restored C57BL/6 (blue) using SplinectomeR. **A)** *Alistipes* (p=0.001), **B)** *Akkermansia* (p=0.001), **C)** *Bacteroides* (p=0.001) **D)** *Erysipelotrichaeae* (p=0.001), **E)** *Anaeroplasma* (p=0.001)

## Notes

**Funding Sources:** This study is supported by research grants from the Masonic Cancer Center ChainBreaker Fund, Mezin Koats colorectal cancer research fund, Minnesota Colorectal Cancer Funds, and research funds from the Department of Surgery and CTSI, University of Minnesota. The Minnesota Colorectal Cancer Research Foundation supports TJG graduate fellowship.

https://www.ncbi.nlm.nih.gov/sra/PRJNA915052

### Competing Interest Statement

The authors have declared no competing interest.

